# How the Body Shapes the Mind’s Eye: Cardiac vagal reactivity predicts visual imagery vividness

**DOI:** 10.64898/2026.05.12.724726

**Authors:** Xindan Zhang, Timo Kvamme, Yoko Nagai, Juha Silvanto

## Abstract

Mental imagery is known to be accompanied by autonomic responses, traditionally viewed as merely downstream consequences of imagery. Recent theoretical work has challenged this view, proposing that mental imagery requires the integration of cortical sensory representations with ascending interoceptive signals supplied by the autonomic nervous system. These two views make opposite predictions: if autonomic activity is only a consequence of imagery, then the responsiveness of the autonomic nervous system should not predict imagery vividness. If instead autonomic input shapes the generation of mental images, individuals with greater autonomic responsiveness should experience more vivid imagery. The present study tested these competing predictions by examining whether individual differences in cardiac vagal reactivity predict self-reported visual imagery vividness, with imagery assessed at a separate time point from the paced breathing protocol. Cardiac vagal reactivity significantly predicted VVIQ scores after controlling for factors such as resting HRV, respiratory amplitude, age and negative affect (b = 13.75, p = .022; model R^2^ = 0.12), whereas neither baseline HRV nor fast-breathing evoked HRV change were linked to imagery, suggesting the effect is specific to the capacity to enter a high parasympathetic state rather than reflecting general autonomic capacity. Self-reported interoceptive noticing (MAIA Noticing) was significantly associated with both delta HRV and VVIQ, suggesting that cardiac vagal reactivity and imagery vividness are linked through a broader individual difference in interoceptive function. These findings indicate that the vividness of mental imagery is not exclusively central in origin but also appears to be shaped by the capacity of the autonomic nervous system to enter a high parasympathetic state. Imagery thus likely involves bidirectional interaction between cortical and autonomic processes, with descending efferent pathways activating autonomic responses while ascending interoceptive signals contributing to the construction of mental images.

## 1. Introduction

Mental imagery (the capacity to generate sensory-like representations in the absence of external stimulation) has been extensively investigated in terms of its neural basis, with neuroimaging studies identifying a range of brain areas and networks involved in imagery generation and maintenance (see e.g. Pearson, 2019; Liu & Bartolomeo, 2025; Spagna et al., 2021). Alongside this cortical literature, early studies established that mental imagery is accompanied by autonomic responses. For example, emotional imagery activates cardiac and other physiological responses resembling those elicited by actual perception, and individual differences in imagery ability modulate the magnitude of these responses. Heart rate acceleration during fearful imagery is greater in individuals with higher imagery ability (Miller et al., 1987), and respiratory responses during high-arousal imagery are likewise more pronounced in higher-ability imagers (e.g. Van Diest et al., 2001; cf. Carroll et al., 1979; Hirschman & Favaro, 1980).

These findings are generally explained in terms of Lang’s bioinformational theory, which proposes that imagery is constructed from associative memory networks containing stimulus, semantic, and response information, with mental imagery automatically recruiting associated autonomic responses via descending efferent pathways (Lang, 1979; Bradley et al., 2023; see also Ji et al, 2016). On this account, autonomic responses are efferent consequences of imagery activation but importantly, do not shape it. In other words, the causal flow is strictly top-down, from cortical image representation to bodily response. Recently, this view has been challenged by the proposal that afferent autonomic input plays a key role in imagery generation. The interoceptive model (Silvanto & Nagai, 2025) proposes that imagery requires the integration of interoceptive signals (conveyed via vagal afferent pathways through the insular cortex) with stored sensory representations, with the anterior insular cortex conferring affective tone, sense of agency, and self-referential grounding to the resulting mental simulation. This account has been elaborated and integrated with attention-based accounts in a dual-stream framework (Scholz et al., 2026).

These two accounts make opposite predictions about the role of the autonomic nervous system in imagery. On the traditional efferent view (Lang, 1979), the functional responsiveness of the autonomic nervous system should be unrelated to imagery vividness; imagery activates autonomic responses via descending efferent pathways, but these responses are a consequence of the mental image that has already been constructed cortically. On the interoceptive account, autonomic input is itself a determinant of imagery, and therefore its properties should shape imagery capacity, with individuals showing greater autonomic responsiveness experiencing more vivid imagery (Silvanto & Nagai, 2025). Prior studies measuring autonomic responses during imagery (e.g. Carroll et al., 1979; Hirschman & Favaro, 1980) cannot distinguish between these accounts, as autonomic activity and imagery occur simultaneously, making it impossible to determine whether autonomic responses are driven by or causal to imagery.

The present study was designed to resolve this ambiguity by measuring autonomic responsiveness temporally separated from imagery assessment, ensuring that any observed relationship between autonomic capacity and imagery vividness cannot reflect autonomic activation as a downstream consequence of imagery. We indexed autonomic responsiveness via heart rate variability (HRV; the beat-to-beat variation in cardiac inter-interval timing) assessed during a paced breathing manipulation. Specifically, we focused on HRV reactivity, i.e. the degree to which HRV increases in response to slow paced breathing, which provides a controlled probe of the capacity to enter a high parasympathetic state (Laborde et al., 2018; Manser et al., 2021). Slow breathing at approximately six breaths per minute corresponds to the resonance frequency of the cardiovascular system, maximally synchronizing respiratory and cardiovascular oscillations and producing robust increases in HRV, with individual differences in the magnitude of this response reflecting stable variation in cardiac vagal capacity across individuals (Sevoz-Couche & Laborde, 2022; Laborde et al., 2022). Importantly for the present study, cardiac vagal control and interoception are closely linked components of a common homeostatic system, with higher vagally-mediated HRV associated with more effective interoceptive and self-regulatory processing (Pinna & Edwards, 2020; Lischke et al., 2021).

We thus examined whether individual differences in cardiac vagal reactivity during slow breathing are associated with visual imagery vividness. Importantly, cardiorespiratory flexibility was assessed at a separate time point from imagery vividness, ensuring that any observed association cannot reflect autonomic activation elicited by imagery itself. On Lang’s (1979) bioinformational account, individual differences in cardiorespiratory flexibility should be unrelated to imagery vividness, since autonomic responses are consequences of cortically constructed images rather than contributors to their generation. On the interoceptive account (Silvanto & Nagai, 2025), by contrast, greater cardiac vagal reactivity should predict more vivid imagery, since individuals who more readily enter a high vagal state engage interoceptive processing more effectively, providing a stronger basis for the bodily signals integrated into mental simulation. Secondly, we examined whether any impact of cardiac vagal reactivity on imagery occurs via interoceptive processing, i.e. processing and detection of bodily signals. Imagery vividness has been shown to relate to both objective and subjective measures of interoception (e.g. Nagai et al., 2026; Monzel et al., 2025; Kvamme et al., 2026), suggesting that the capacity to detect and represent bodily signals is a consistent correlate of imagery ability. To test whether any autonomic association with imagery vividness is related to interoceptive processing, we also assessed subjective interoceptive awareness (MAIA) and examined whether it accounts for the relationship between autonomic reactivity and imagery vividness.

## 2. Methods

### 2.1. Participants

A total of 112 participants with normal or corrected-to-normal vision were recruited from the University of Macau. Written informed consent was obtained from all participants prior to testing, and participants received financial compensation upon completion. After excluding participants with excessive physiological noise, incomplete recordings, or values exceeding the ±3 SD outlier criterion, the final analytic sample consisted of 108 participants (mean age = 23.92 years, SD = 2.86; 84 female participants, 24 male participants).The study received ethical approval from the University of Macau Research Ethics Committee and was conducted in accordance with the principles of the Declaration of Helsinki.

### 2.2. Experimental Design

The study employed a within-subjects repeated-measures design. Each participant completed three 5-minute physiological conditions: resting state, slow-paced breathing at 6 breaths per minute, and fast-paced breathing at 20 breaths per minute. During the paced-breathing conditions, participants followed a visual breathing guide presented on the screen. A white rectangle moved rhythmically upward and downward, and the words “inhale” and “exhale” appeared on the screen during the corresponding inspiratory and expiratory phases. In the slow-breathing condition, each inhalation and exhalation phase lasted 5 s, yielding a fixed respiratory frequency of 0.10 Hz (6 breaths/min). In the fast-breathing condition, each inhalation and exhalation phase lasted 1.5 s, yielding a fixed respiratory frequency of 0.33 Hz (20 breaths/min). Because all participants followed the same externally paced visual waveform, respiratory frequency was standardized across individuals within each condition. Compliance with the paced breathing was monitored in real time using the AcqKnowledge platform to ensure that each participant maintained the prescribed respiratory frequency throughout the recording. Respiratory depth was quantified from the respiratory belt signal and statistically controlled in subsequent analyses. The resting-state condition was always administered first to obtain a baseline measure of cardiac variability before paced-breathing manipulation. During this condition, participants were instructed to close their eyes, remain physically relaxed, and rest quietly for 5 minutes. The order of the slow- and fast-paced breathing conditions was counterbalanced across participants.

The questionnaire battery included the Vividness of Visual Imagery Questionnaire Chinese version (VVIQ-C; Marks, 1973; Zhang et al., 2025) as the primary outcome measure, and the Multidimensional Assessment of Interoceptive Awareness (MAIA-2C; Mehling et al., 2018; Teng et al., 2022) as the **primary subjective interoceptive measure**. The Generalized Anxiety Disorder-7 scale (GAD-7; Spitzer et al., 2006; He et al., 2010), the Patient Health Questionnaire-9 (PHQ-9; Kroenke et al., 2001; Wang et al., 2014), and the State–Trait Anxiety Inventory trait subscale (STAI-trait; Du et al., 2022) were included as control measures to account for the potential confounding influence of negative affect on the association between cardiac vagal reactivity and imagery vividness.

To reduce the possibility that questionnaire responses reflected transient effects of the breathing protocol, the order of the physiological protocol and questionnaire battery was varied across participants. Specifically, 56 participants completed the paced-breathing protocol before the questionnaire battery, whereas 56 participants completed the questionnaire battery before the paced-breathing protocol. A 10-minute break was inserted between the two parts of the experiment. Thus, participants who completed the breathing protocol first rested for 10 minutes before completing the questionnaire battery, whereas participants who completed the questionnaire battery first rested for 10 minutes before beginning the breathing protocol. HRV has been shown to return to baseline immediately following cessation of slow breathing, with RMSSD showing no significant difference between pre- and post-breathing measurements after a 5-minute session (Laborde et al., 2022). The 10-minute rest period was therefore a conservative buffer, sufficient to ensure that questionnaire responses reflect stable trait-level imagery ability rather than any transient breathing-induced state.

### 2.3. Physiological Data Acquisition

Continuous ECG and respiratory signals were recorded at 2000 Hz using a BioPac MP160 system with AcqKnowledge acquisition software. Raw ECG data were preprocessed to remove baseline drift using a zero-phase second-order Butterworth high-pass filter with a cutoff frequency of 0.5 Hz. QRS complexes were subsequently enhanced via discrete wavelet decomposition using a Symlet-4 wavelet with six decomposition levels. Signal reconstruction was based on a selective combination of detail coefficients from decomposition levels 4, 5, and 6, weighted at 0.3, 1.0, and 1.5, respectively. R-peaks were then localized using an adaptive detection algorithm, with a minimum inter-peak distance of 0.35 times the sampling frequency and a minimum peak prominence threshold set at 30% of the amplitude range between the 80th and 97th percentiles of the enhanced signal.

Respiratory recordings were linearly detrended and used to verify breathing compliance and quantify respiratory amplitude. Respiratory rate was extracted from the recorded respiratory belt signal to confirm that participants followed the paced-breathing instructions. Respiratory amplitude was estimated from the respiratory belt signal as the peak-to-trough excursion of each respiratory cycle. Participants were excluded if physiological recordings showed substantial ECG data loss over 30%, or insufficient valid R–R intervals for HRV estimation. Univariate outliers on HRV indices were identified using a criterion of ±3 SD from the sample mean.

### 2.4. Time-Domain HRV and Respiratory-Frequency Spectral Measures

Heart rate variability was quantified using the root mean square of successive differences between consecutive normal R–R intervals (RMSSD), a standard time-domain measure of short-term cardiac variability. Mean heart period (mean R–R interval, ms) was computed for each condition to allow joint examination of HP and HRV per Quigley et al. (2024) recommendations. Because RMSSD can depend mathematically on mean heart period, RMSSD values were normalized by the mean R–R interval for each condition (RMSSD/ mean R–R interval).

In addition to time-domain RMSSD, frequency-domain HRV indices were computed to characterize cardiac oscillatory power at the paced respiratory frequency. For the slow-paced breathing condition, heart-period power was extracted from a narrow band centered on the target breathing frequency of 0.10 Hz, spanning 0.07–0.13 Hz. For the fast-paced breathing condition, heart-period power was extracted from an analogous band centered on the target breathing frequency of 0.33 Hz, spanning 0.23–0.43 Hz. These bands corresponded to approximately ±30% around each target breathing frequency and were selected to capture respiratory-frequency cardiac oscillatory activity while accommodating small deviations in actual breathing timing and spectral leakage inherent in finite-length physiological recordings. These measures are referred to as slow-breathing 0.10-Hz HRV power and fast-breathing 0.33-Hz HRV power, respectively.

### 2.5. Statistical Analyses

All analyses were conducted in R. VVIQ scores were standardized prior to regression analyses. Pearson correlations were first used to examine zero-order associations between VVIQ scores and HRV indices across breathing conditions, including resting RMSSD, slow-breathing RMSSD, and fast-breathing RMSSD. The primary analysis tested whether individual differences in slow-breathing–evoked cardiac vagal reactivity were associated with imagery vividness. Cardiac vagal reactivity was operationalized as the change in RMSSD normalized by mean R–R interval from rest to slow-paced breathing. A linear regression model was fitted with standardized VVIQ scores as the dependent variable and slow-breathing–evoked RMSSD change as the primary independent variable. The model included resting RMSSD normalized by mean R–R interval, respiratory amplitude change from rest to slow breathing, resting respiratory amplitude, age, and a standardized negative-affect composite as covariates. The negative-affect composite was created by z-standardizing and averaging GAD-7, PHQ-9, and STAI-trait scores. This composite was included to test whether the association between slow-breathing–evoked RMSSD change and VVIQ scores could be explained by general anxiety symptom burden or trait anxiety.

A convergent frequency-domain regression model was then fitted to test whether slow-breathing 0.1-Hz HRV power predicted VVIQ. This model included slow-breathing 0.1-Hz HRV power as the primary predictor and fast-breathing 0.33-Hz HRV power, respiratory amplitude of slow and fast breathing state, negative-affect composite, and age as covariates. This analysis tested whether imagery vividness was associated with respiratory-frequency cardiac oscillatory power during the 0.1-Hz slow-breathing challenge.

To examine whether the association between cardiac vagal reactivity and imagery vividness is statistically accounted for by subjective interoceptive awareness, an exploratory bootstrapped mediation analysis was conducted. MAIA Noticing was selected as the mediator because it was the only MAIA subscale that correlated significantly with both delta HRV and VVIQ (see Results), making it the candidate interoceptive variable for this analysis. Delta HRV was entered as the independent variable, VVIQ as the outcome, and MAIA Noticing as the mediator. Indirect, direct, and total effects were estimated using 5,000 bootstrap resamples with bias-corrected 95% confidence intervals.

## 3. Results

### 3.1. Descriptive statistics

Descriptive statistics for all autonomic indices and imagery vividness are summarized in Table 1. VVIQ scores were normally distributed (mean = 65.21, SD = 9.51) and covered a wide range of imagery vividness (range: 40–80), confirming adequate variability for correlational analyses.

**Table 1.**
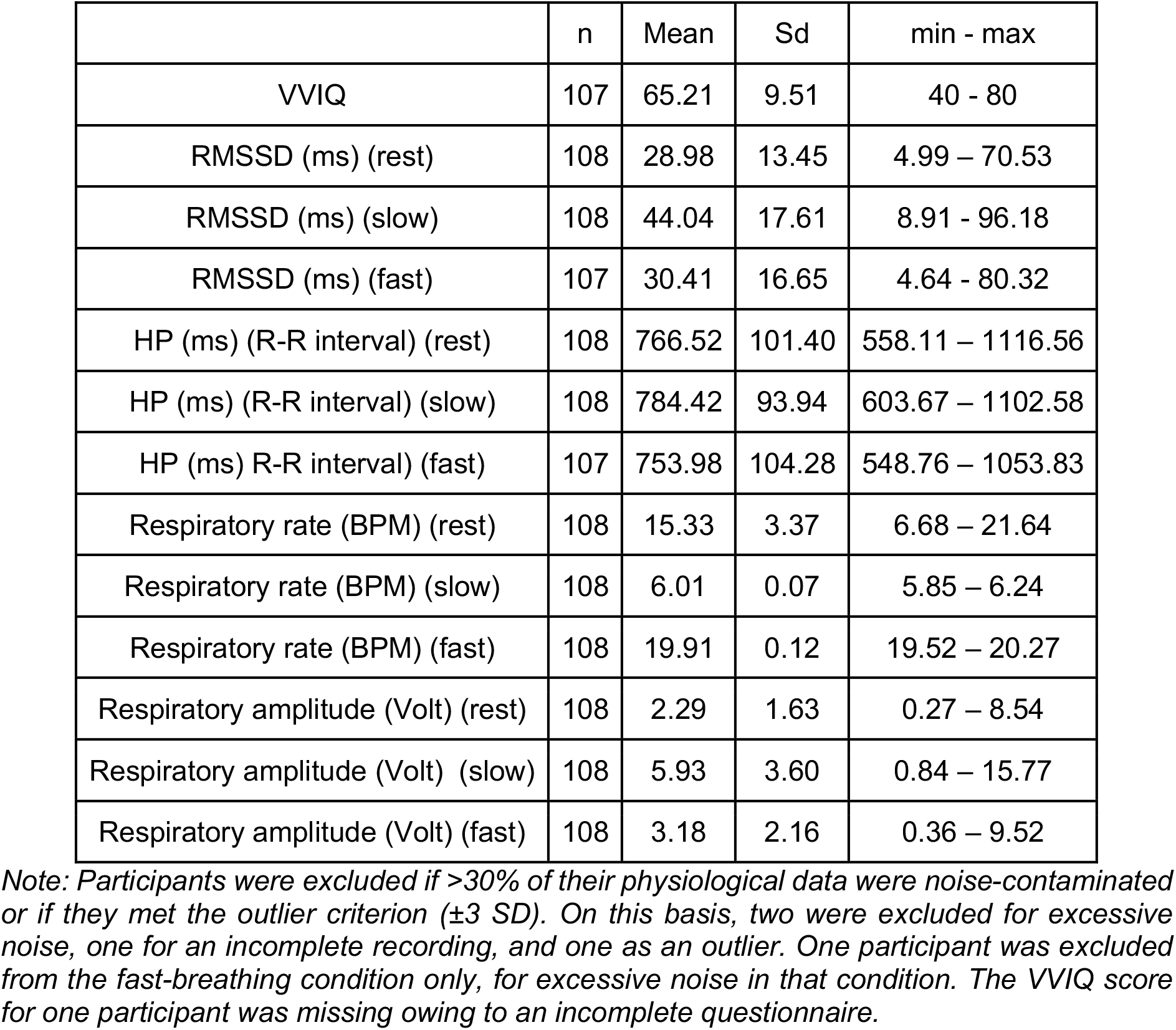
Descriptive Statistics.

### 3.2. Paced Breathing Modulated Cardiorespiratory Dynamics

To verify that the paced-breathing manipulation altered respiratory and cardiac dynamics, heart period (HP; R–R interval, ms), RMSSD (ms), and respiratory amplitude were compared across slow-breathing, resting, and fast-breathing conditions using repeated-measures ANOVAs. Significant main effects of breathing condition were observed for HP, *F*(2, 212) = 16.47, *p* < .001, RMSSD, *F*(2, 212) = 95.84, *p* < .001, and respiratory amplitude, *F*(2, 214) = 135.80, *p* < .001.

Bonferroni-corrected paired-samples *t* tests showed that HP was significantly higher during slow breathing than during both resting (*t*(106) = 3.53, *p* < .001, mean difference = 17.97 ms, 95% CI [7.88, 28.06]), and fast breathing (*t*(106) = 5.88, *p* < .001, mean difference = 28.35 ms, 95% CI [18.80, 37.90]). And the HP was significantly higher during resting breathing than during fast breathing, *t*(106) = 2.04, *p* = .04). For RMSSD, values were significantly higher during slow breathing than during both resting (*t*(106) = 11.69, *p* < .001, mean difference = 15.22 ms, 95% CI [12.64, 17.80]), and fast breathing (*t*(106) = 10.65, *p* < .001, mean difference = 13.64 ms, 95% CI [11.10, 16.18]). RMSSD did not differ significantly between resting and fast breathing (*t*(106) = −1.55, *p* = .13). Respiratory amplitude also differed significantly across conditions. Follow-up comparisons showed that respiratory amplitude was significantly higher during slow breathing than during both resting (*t*(107) = 12.28, *p* < .001, mean difference = 3.64, 95% CI [3.05, 4.23]), and fast breathing (*t*(107) = 12.95, *p* < .001, mean difference = 2.75, 95% CI [2.33, 3.17]). Respiratory amplitude was also significantly higher during fast breathing than during resting (*t*(107) = 5.52, *p* < .001, mean difference = 0.89, 95% CI [0.57, 1.21]).

Together, these findings indicate that the paced-breathing manipulation successfully altered respiratory amplitude and cardiac variability. During slow-paced breathing, heart period increased and RMSSD increased relative to resting and fast-breathing conditions. This pattern is consistent with physiological signatures of vagally mediated cardiac regulation, indicating that the manipulation effectively engaged cardiorespiratory processes associated with parasympathetic modulation.

### 3.3. Slow-Breathing–Evoked cardiac vagal reactivity Is associated with Imagery Vividness

We first examined zero-order correlations between VVIQ scores and HRV indices across the breathing condition. These are shown in Table 2. VVIQ scores were positively correlated with the slow-minus-rest RMSSD change score (r = .229, p = .018; see Figure 1A). In contrast, VVIQ scores were not significantly associated with RMSSD during rest (r = .045, p = .643) or fast-minus-rest RMSSD change score (r = .006, p = .948; see Figure 1B). This pattern suggests that imagery vividness was most consistently related to cardiac variability expressed during, and in response to, the slow-breathing condition.

**Table 2.**
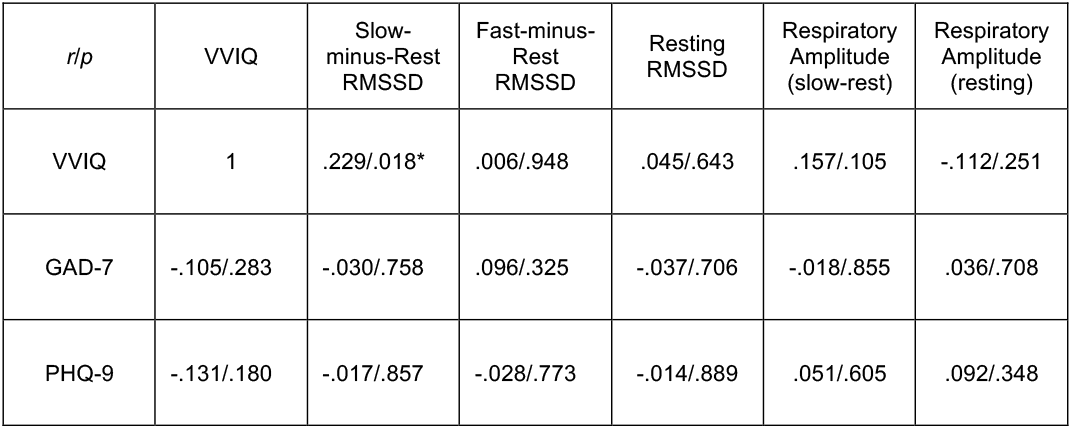
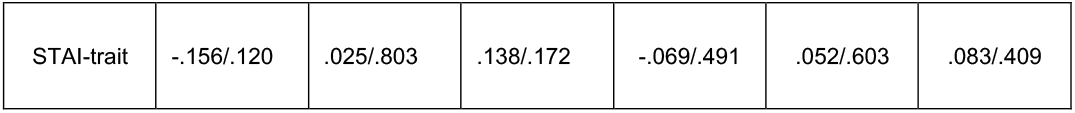
Correlations (Pearson’s r / p-value) between Questionnaire measures and HRV

**Figure 1.**
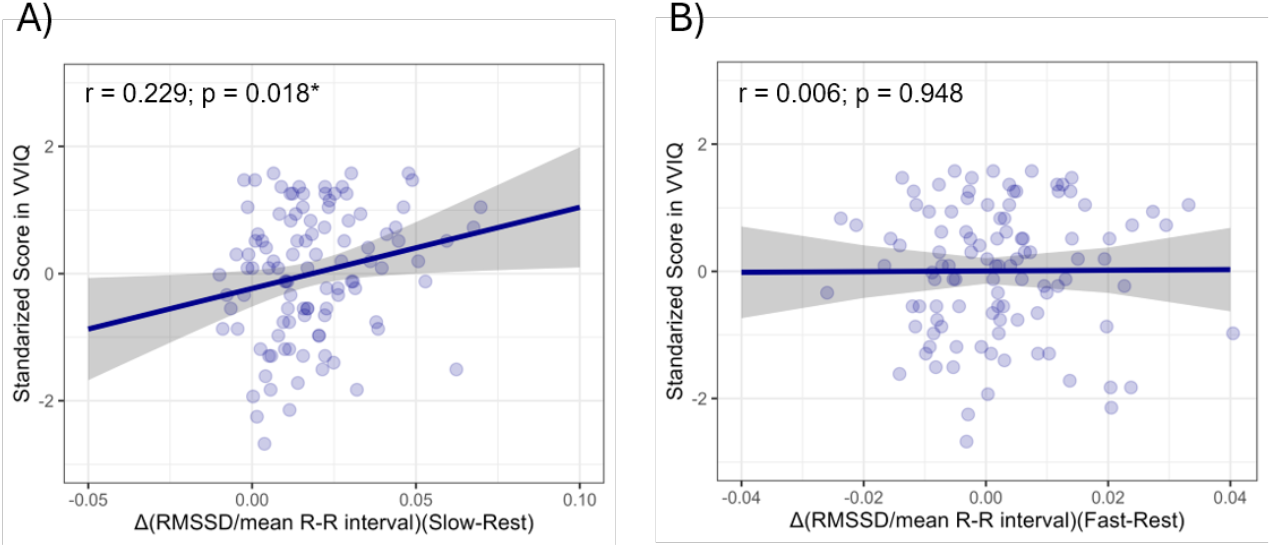
Cardiac variability during paced breathing and visual imagery vividness. Scatter plots show associations between VVIQ scores and HRV indices across breathing conditions. (A) Slow-breathing–evoked RMSSD change, normalized by mean R–R interval, was significantly associated with VVIQ scores. (B) Fast-breathing–evoked RMSSD change, normalized by mean R–R interval, was not significantly associated with VVIQ scores. Shaded regions indicate 95% confidence intervals.

We then fitted the primary baseline-adjusted regression model described in section 2.5, which regressed standardized VVIQ scores onto slow-breathing–evoked RMSSD change, controlling for resting RMSSD, respiratory amplitude change, resting respiratory amplitude, age, and a negative affect composite comprising anxiety and depression measures (GAD-7, PHQ-9, and STAI-trait). The overall model was significant (F(6, 100) = 2.214, p = .048, R^2^ = .117). Slow-breathing–evoked RMSSD change significantly predicted VVIQ scores (b = 13.747, SE = 5.905, t(100) = 2.328, p = .022). And the remaining covariates were not significant predictors. Because respiratory amplitude was statistically controlled and the primary effect survived, it can be concluded that the slow-breathing–evoked RMSSD effect on VVIQ partly reflects cardiac vagal reactivity rather than purely respiratory mechanisms.

To determine whether this association extended to frequency-domain cardiac variability, we fitted the convergent frequency-domain regression models described in Section 2.5. Specifically, we examined heart-period power centered on the paced slow-breathing frequency of 0.10 Hz and the paced fast-breathing frequency of 0.33 Hz. Slow-breathing 0.10- Hz HRV power positively predicted VVIQ scores (b = 0.362, SE = 0.172, t(99) = 2.101, p = .038, R^2^ = .092). In the same model, fast-breathing 0.33-Hz HRV power did not independently predict VVIQ scores (b = -0.106, SE = 0.107, t(99) = -0.991, p = .324). These findings suggest that imagery vividness was associated not only with time-domain RMSSD responsiveness during slow breathing, but also with respiratory-frequency cardiac oscillatory power during the 0.10-Hz slow-breathing challenge.

In summary, greater RMSSD increases from rest to slow-paced breathing were associated with higher visual imagery vividness. This result suggests that individuals with greater cardiorespiratory flexibility during a 0.1-Hz slow-breathing challenge tended to report more vivid mental imagery.

### 3.4. The relationship between Self-reported interoception (MAIA), HRV and VVIQ

We next examined whether imagery vividness was related to self-reported interoceptive awareness. The correlation matrix is provided in Table 3. MAIA Noticing subscale was positively correlated with VVIQ (r = .450, p <.001) and slow-minus-rest HRV (r = .200, p = .039). While Emotion Awareness (r = .343 p <.001), and Trust subscales (r = .241, p = .013) were significantly correlated with VVIQ, they were not with HRV measures.

**Table 3.** Relationship between MAIA, VVIQ and HRV

| *r/p* | VVIQ | Slow-minus-Rest<br>RMSSD | Resting RMSSD |
| --- | --- | --- | --- |
| MAIA (Noticing) | .450/<.001*** | .200/.039* | -.097/.321 |
| MAIA (Not-Distracting) | .015/.882 | -.018/.852 | .012/.904 |
| MAIA (Not-Worrying) | .065/.509 | -.003/.979 | -.010/.92 |
| MAIA (Attention Regulation) | .204/.036* | -.061/.533 | -.007/.939 |
| MAIA (Emotional Awareness) | .343/<.001*** | .144/.139 | .0004/.997 |
| MAIA (Self-Regulation) | .033/.738 | -.046/.640 | .070/.471 |
| MAIA (Body Listening) | .133/.175 | .183/.059 | -.049/.618 |
| MAIA (Trust) | .241/.013* | -.022/.820 | .152/.119 |

Given that MAIA Noticing correlated significantly with both cardiac vagal reactivity (i.e. Slow- minus-Rest RMSSD) and VVIQ, we carried out an exploratory analysis to test whether it mediated the relationship between the two.

A bootstrapped mediation analysis with 5,000 resamples was conducted to test whether interoceptive noticing mediated the relationship between cardiac vagal reactivity (delta HRV) and imagery vividness (VVIQ). Delta HRV significantly predicted MAIA Noticing (path a: b = 0.203, SE = 0.091, 95% CI [0.021, 0.380], z = 2.221, p = .026). MAIA Noticing also significantly predicted VVIQ when controlling for delta HRV (path b: b = 0.421, SE = 0.087, 95% CI [0.254, 0.588], z = 4.857, p < .001). The total effect of delta HRV on VVIQ was significant (c: b = 0.229, SE = 0.091, 95% CI [0.046, 0.405], z = 2.526, p = .012), whereas the direct effect was not significant after including MAIA Noticing in the model (c′: b = 0.144, SE = 0.081, 95% CI [−0.016, 0.303], z = 1.774, p = .076). The indirect effect was significant (a × b = 0.086, SE = 0.040, 95% CI [0.009, 0.167], z = 2.113, p = .035), consistent with an indirect association between delta HRV and VVIQ via MAIA Noticing. In summary, these results indicate that, once interoceptive noticing was included as a mediator, the direct association between cardiac vagal reactivity and imagery vividness was reduced and no longer reached significance (c′ = 0.144, p = .076), while the indirect path through noticing was significant and accounted for approximately 38% of the total effect. This is consistent with partial mediation, though the cross-sectional design does not permit a directed causal interpretation.

## 4. Discussion

The present study tested two opposing accounts of the relationship between the autonomic nervous system and mental imagery. On the efferent view (Lang, 1979), autonomic responses are downstream consequences of cortically constructed images, so individual differences in autonomic capacity should be unrelated to imagery vividness. On the interoceptive account (Silvanto & Nagai, 2025), ascending autonomic signals play a causal role in imagery generation, so individuals with greater cardiac vagal reactivity should report more vivid imagery. Because cardiac vagal reactivity and imagery vividness were assessed at separate time points, any association between them cannot reflect autonomic activation elicited by imagery itself, allowing the two accounts to be distinguished. The results support the interoceptive account, as greater slow-breathing–evoked increases in RMSSD predicted higher VVIQ scores, and this association held after controlling for resting HRV, respiratory amplitude, negative affect, and age. Importantly, because respiratory rate was standardised across participants by the paced-breathing protocol and respiratory amplitude was statistically controlled, individual differences in slow-breathing–evoked RMSSD change cannot be merely attributed to differences in breathing depth. This allows the effect to be interpreted as reflecting cardiac vagal reactivity rather than simply respiratory mechanics, consistent with established practice in paced-breathing HRV research (Laborde et al., 2022; Quigley et al., 2024). In contrast, resting HRV and fast-breathing HRV change did not significantly predict imagery vividness, suggesting the association was specific to the capacity to enter a high vagal state during slow breathing rather than reflecting tonic vagal tone or general autonomic capacity, and argues against a generic autonomic arousal account in which any increase in autonomic activity is linked to imagery. Convergent frequency-domain analyses showed that cardiac oscillatory power at the 0.1-Hz slow-breathing frequency also predicted imagery vividness, providing independent support for the primary RMSSD finding.

The principal implication of these findings is that imagery vividness is not exclusively cortical in origin but also relates to autonomic-interoceptive function. Of the MAIA subscales, Noticing was the only one associated with both delta HRV (r = .20) and VVIQ (r = .45), and the direct HRV–VVIQ path fell to non-significance once it was included in the exploratory mediation analysis. We interpret this within the interoceptive model of imagery (Silvanto & Nagai, 2025), in which imagery generation requires the intergration of ascending visceral signals with stored sensory representations at cortical level. In this view, the capacity to enter a high vagal state predicts imagery vividness because it is associated with more effective interoceptive processing (Pinna & Edwards, 2020; Lischke et al., 2021), which the model holds to support this integration (Silvanto & Nagai, 2025). The reported mediation is best read as showing that the autonomic and interoceptive correlates of imagery overlap, consistent with a shared autonomic-interoceptive disposition across vagal reactivity, interoceptive noticing, and imagery vividness. We cannot interpret it as a direct causal pathway, as the cross-sectional design cannot separate a mediating role for noticing from a common disposition underlying all three, and self-reported noticing is best understood as a correlated marker of this disposition rather than the conscious readout of afferent signal strength, from which it is known to dissociate (Garfinkel et al., 2015).

The anterior insular cortex is the most likely neural substrate of the observed effects. The insula receives continuous vagal afferent input via the nucleus tractus solitarius and parabrachial nucleus, and functions as the primary hub where ascending interoceptive signals converge (e.g. Critchley & Harrison, 2013). According to the interoceptive model of imagery (Silvanto & Nagai, 2025), the anterior insula coordinates the integration of these ascending signals with stored cortical sensory representations, and this integration contributes to the vividness of the resulting imagery. Consistent with this, recent neuroimaging evidence supports the insula’s involvement in mental imagery (Kvamme et al., 2025; Takamura et al., 2026).

The magnitude of the present association is comparable to effect sizes reported in previous studies linking VVIQ to neural, structural, and interoceptive measures. The zero-order association between VVIQ and slow-breathing–evoked RMSSD change (r = .229), and the baseline-adjusted regression effect (R^2^ = .117), are similar in size to reported associations between VVIQ and hippocampal volume (r = .35, R^2^ = .12; Tabi et al., 2022), frontal delta EEG power under working-memory load (r = .30, R^2^ = .09; Boere et al., 2025), cardioceptive sensitivity (Nagai et al., 2026), and self-reported interoceptive measures (Kvamme et al., 2026). Thus, the present HRV effect is within the range typically observed for individual-difference predictors of imagery vividness, supporting the relevance of cardiac vagal reactivity to imagery vividness alongside established neural and structural measures.

From a theoretical perspective, these findings do not support the view that autonomic activation is merely a consequence of vivid imagery. On the traditional efferent account (Lang, 1979), the autonomic nervous system is activated in response to cortically constructed mental image but plays no role in imagery generation. From this it follows that individual differences in autonomic capacity should not predict imagery vividness. In other words, a person with high or low cardiac vagal capacity should image equally vividly, since vividness is determined by cortical processes. Rather, these findings are more parsimoniously accounted for by the interoceptive model (Silvanto & Nagai, 2025) and its elaboration in the dual-stream framework (Scholz et al., 2026), both of which propose that interoceptive signals play a causal role in imagery generation. A parsimonious view incorporating both theories is that imagery likely involves bidirectional interaction between cortical and autonomic processes, with descending efferent pathways activating autonomic responses while ascending interoceptive signals contributing to the construction of mental images.

Several alternative explanations for the present findings can be ruled out. One concern is that a stable third variable (such as general physiological fitness) could independently drive both higher vagal reactivity and more vivid imagery, producing the observed association without any direct link between the two. A related concern is that self-reported imagery vividness may be influenced by response biases such as social desirability, or that the finding reflects general cognitive capacity rather than imagery vividness specifically. Multiple aspects of the results argue against all of these interpretations. Firstly, the relationship of VVIQ and delta HRV to MAIA noticing subscale points to a genuine individual difference in the interoceptive-autonomic architecture supporting mental simulation rather than to a generic confound. Secondly, the primary regression controlled for a negative affect composite comprising anxiety and depression measures. Thirdly, resting HRV, a well-established correlate of both general psychological wellbeing and executive function (Thayer & Lane, 2000; Forte et al., 2019) did not predict imagery vividness, indicating that the effect is specific to dynamic vagal reactivity rather than tonic autonomic health or general cognitive capacity.

Certain limitations of the present study nonetheless need to be acknowledged. The present results cannot establish that cardiorespiratory flexibility causally determines imagery vividness; experimental manipulations of bodily or cardiorespiratory state during mental imagery will be necessary to test this possibility directly. In the present study, paced breathing and imagery were examined separately in order to test the specific hypothesis on the link between autonomic reactivity and imagery vividness. Had imagery assessment been done during paced breathing, it would have been impossible to adjudicate between the two theories. A further limitation is that the sample was drawn from a university student population with a restricted age range and a primarily female composition. Future studies are needed to examine whether the relationship between imagery vividness and autonomic-interoceptive functioning varies across the lifespan, across genders, and in more diverse populations.

In conclusion, the present study provides physiological evidence that cardiac vagal capacity predicts visual imagery vividness. Notably, resting HRV did not predict imagery vividness, indicating that the effect is specific to dynamic vagal reactivity rather than reflecting a general physiological fitness confound. Rather than a unidirectional relationship, imagery may require bidirectional interaction between cortical and autonomic processes, with descending efferent pathways activating autonomic responses while ascending interoceptive signals contribute to the construction of mental images. The vividness of mental imagery is thus not exclusively central in origin; it is also shaped peripherally, by the capacity of the autonomic nervous system to contribute to the interoceptive basis of embodied simulation.

## Funding acknowledgements

This work was funded by grants from University of Macau (SRG2025-00010-ICI) and the BIAL Foundation (25/24) awarded to JS.

## Data availability

All data is publicly available at: https://osf.io/97s3n/overview

---

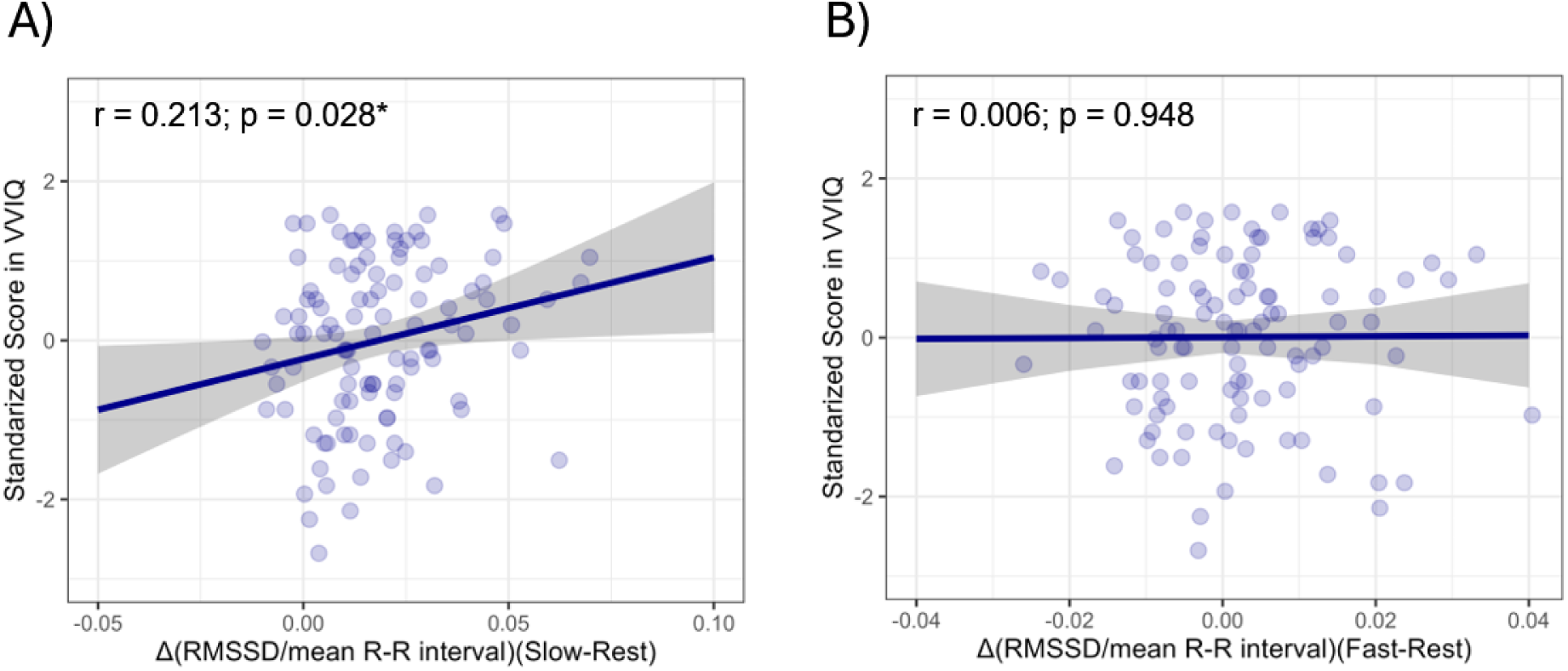

## References

Bradley, M. M., Sambuco, N., & Lang, P. J. (2023). Imagery, emotion, and bioinformational theory: From body to brain. Biological Psychology, 183, 108669.

Boere, K., Krempel, R., Walsh, E., Li, H., McLaughlin, L., Krigolson, O. E., & Blomkvist, A. (2025). Task evoked EEG reveals neural processing differences in aphantasia. Scientific Reports.

Carroll, D., Baker, J., & Preston, M. (1979). Individual differences in visual imaging and the voluntary control of heart rate. British Journal of Psychology, 70(1), 39–49.

Craig, A. D. (2009). How do you feel — now? The anterior insula and human awareness. Nature Reviews Neuroscience, 10(1), 59–70.

Critchley, H. D. (2005). Neural mechanisms of autonomic, affective, and cognitive integration. Journal of Comparative Neurology, 493, 154–166.

Critchley, H. D., & Harrison, N. A. (2013). Visceral influences on brain and behavior. Neuron, 77(4), 624–638.

Du, Q., Liu, H., Yang, C., Chen, X., & Zhang, X. (2022). The development of a short Chinese version of the state-trait anxiety inventory. Frontiers in Psychiatry, 13, 854547.

Forte, G., Favieri, F., & Casagrande, M. (2019). Heart rate variability and cognitive function: A systematic review. Frontiers in Neuroscience, 13, 710.

Garfinkel, S.N., Seth, A.K., Barrett, A.B., Suzuki, K., Critchley, H.D. (2015). Knowing your own heart: distinguishing interoceptive accuracy from interoceptive awareness. Biol Psychol. Jan;104:65–74. doi: 10.1016/j.biopsycho.2014.11.004.

Hirschman, R., & Favaro, L. (1980). Individual differences in imagery vividness and voluntary heart rate control. Personality and Individual Differences, 1(2), 129–133.

He, X. Y. L. C., Li, C., Qian, J., Cui, H. S., & Wu, W. Y. (2010). Reliability and validity of a generalized anxiety disorder scale in general hospital outpatients. Shanghai Arch Psychiatry, 22(4), 200–203.

Ji, J. L., Heyes, S. B., MacLeod, C., & Holmes, E. A. (2016). Emotional mental imagery as simulation of reality: Fear and beyond—A tribute to Peter Lang. Behavior Therapy, 47(5), 702–719.

Kvamme, T. L., Lumaca, M., Bajada, C. J., Gregersen, S. D., Hobot, J., Paunovic, D., Wierzchon, M., Zana, B., Silvanto, J., & Sandberg, K. (2025). Neural network topologies supporting individual variations in vividness of visual imagery. NeuroImage, 321, 121520.

Kvamme, T. L., Monzel, M., Nagai, Y., & Silvanto, J. (2026). When weak imagery is worse than none: Core aphantasia and hypophantasia relate differently to mental health, mediated by subjective interoception. Neuropsychologia, 222, 109368.

Kroenke, K., Spitzer, R. L., & Williams, J. B. (2001). The PHQ-9: validity of a brief depression severity measure. Journal of general internal medicine, 16(9), 606–613.

Laborde, S., Allen, M. S., Borges, U., Dosseville, F., Hosang, T. J., Iskra, M., & Javelle, F. (2022). Effects of voluntary slow breathing on heart rate and heart rate variability: A systematic review and a meta-analysis. Neuroscience & Biobehavioral Reviews, 138, 104711.

Laborde, S., Mosley, E., & Mertgen, A. (2018). Vagal tank theory: the three rs of cardiac vagal control functioning–resting, reactivity, and recovery. Frontiers in neuroscience, 12, 458.

Lang, P. J. (1979). A bio-informational theory of emotional imagery. Psychophysiology, 16(6), 495–512.

Lischke. A., Pahnke, R., Mau-Moeller, A. & Weippert. M. (2021). Heart Rate Variability Modulates Interoceptive Accuracy. Front. Neurosci. 14:612445. doi: 10.3389/fnins.2020.612445

Liu, J., & Bartolomeo, P. (2025). Aphantasia as a functional disconnection. Trends in Cognitive Sciences, 29(11), 963–964.

Marks, D. F. (1973). Visual imagery differences in the recall of pictures. British journal of Psychology, 64(1), 17–24.

Miller, G. A., Levin, D. N., Kozak, M. J., Cook III, E. W., McLean Jr, A., & Lang, P. J. (1987). Individual differences in imagery and the psychophysiology of emotion. Cognition and emotion, 1(4), 367–390.

Manser, P., Thalmann, M., Adcock, M., Knols, R. H., & de Bruin, E. D. (2021). Can reactivity of heart rate variability be a potential biomarker and monitoring tool to promote healthy aging? A systematic review with meta-analyses. Frontiers in Physiology, 12, 686129.

Mehling, W. E., Acree, M., Stewart, A., Silas, J., & Jones, A. (2018). The multidimensional assessment of interoceptive awareness, version 2 (MAIA-2). PloS one, 13(12), e0208034.

Monzel, M., Nagai, Y., & Silvanto, J. (2025). The role of subjective interoception in autobiographical deficits in aphantasia. Scientific Reports, 15, 43007.

Nagai, Y., Arooj, S., Futeran-Blake, T. R., Manders, C., Critchley, H., & Silvanto, J. (2026). Interoception predicts mental imagery vividness: exploring a key relationship. Scientific Reports, 16, 14181. 10.1038/s41598-026-43805-0

Pearson, J. (2019). The human imagination: the cognitive neuroscience of visual mental imagery. Nature Reviews Neuroscience, 20(10), 624–634.

Pinna, T., Edwards, D.J. (2020) A Systematic Review of Associations Between Interoception, Vagal Tone, and Emotional Regulation: Potential Applications for Mental Health, Wellbeing, Psychological Flexibility, and Chronic Conditions. Front. Psychol. 11:1792. doi: 10.3389/fpsyg.2020.01792

Quigley, K. S., Gianaros, P. J., Norman, G. J., Jennings, J. R., Berntson, G. G., & de Geus, E. J. (2024). Publication guidelines for human heart rate and heart rate variability studies in psychophysiology—Part 1: Physiological underpinnings and foundations of measurement. Psychophysiology, 61(9), e14604.

Spitzer, R. L., Kroenke, K., Williams, J. B., & Löwe, B. (2006). A brief measure for assessing generalized anxiety disorder: the GAD-7. Archives of internal medicine, 166(10), 1092–1097.

Scholz, C., Monzel, M., Kvamme, T., Liu, J., & Silvanto, J. (2026). An Integration Model of Mental Imagery and Aphantasia: Conceptual Framework, Neuromechanistic Pathways, and Clinical Implications. Neuropsychologia 225:109401.

Sevoz-Couche, C., & Laborde, S. (2022). Heart rate variability and slow-paced breathing: when coherence meets resonance. Neuroscience & Biobehavioral Reviews, 135, 104576.

Silvanto, J., & Nagai, Y. (2025). How interoception and the insula shape mental imagery and aphantasia. Brain Topography, 38(2), 27.

Spagna, A., Hajhajate, D., Liu, J., & Bartolomeo, P. (2021). Visual mental imagery engages the left fusiform gyrus, but not the early visual cortex: A meta-analysis of neuroimaging evidence. Neuroscience & Biobehavioral Reviews, 122, 201–217.

Teng, B., Wang, D., Su, C., Zhou, H., Wang, T., Mehling, W. E., & Hu, Y. (2022). The multidimensional assessment of interoceptive awareness, version 2: Translation and psychometric properties of the Chinese version. Frontiers in psychiatry, 13, 970982.

Tabi, Y. A., Maio, M. R., Attaallah, B., Dickson, S., Drew, D., Idris, M. I., Kienast, A., Klar, V., Nobis, L., Plant, O., Saleh, Y., Sandhu, T. R., Slavkova, E., Toniolo, S., Zokaei, N., Manohar, S. G., & Husain, M. (2022). Vividness of visual imagery questionnaire scores and their relationship to visual short-term memory performance. Cortex, 146, 186–199.

Takamura, Y., Delsanti, R., Cohen, L., Bartolomeo, P., & Liu, J. (2026). Congenital aphantasia reveals frontotemporal and cingulate structural alterations underlying conscious access to imagery. bioRxiv, 2026–03.

Thayer, J. F., & Lane, R. D. (2000). A model of neurovisceral integration in emotion regulation and dysregulation. Journal of Affective Disorders, 61(3), 201–216.

Van Diest, I., Winters, W., Devriese, S., Vercamst, E., Han, J. N., Van de Woestijne, K. P., & Van den Bergh, O. (2001). Hyperventilation beyond fight/flight: respiratory responses during emotional imagery. Psychophysiology, 38(6), 961–968.

Wang, W., Bian, Q., Zhao, Y., Li, X., Wang, W., Du, J., … & Zhao, M. (2014). Reliability and validity of the Chinese version of the Patient Health Questionnaire (PHQ-9) in the general population. General hospital psychiatry, 36(5), 539–544.

Zhang, Z., Liu, Y., Yang, J., Li, C., Marks, D. F., Della Sala, S., & Zhao, B. (2025). The establishment of the Chinese version of the Vviq (Vviq-C) and the effects of age and gender on the vividness of visual imagery. Journal of Psychiatry and Psychiatric Disorders, 9(3), 163–171.

